# Delivery of human adipose-derived stromal cells within *in situ* forming chitosan/PEG-PTMC hydrogels induces adverse outcomes in a femoral artery ligation model in athymic *nu/nu* mice

**DOI:** 10.1101/2023.05.03.539275

**Authors:** Fiona E. Serack, John A. Ronald, Brian G. Amsden, David A. Hess, Lauren E. Flynn

**Affiliations:** School of Biomedical Engineering, Faculty of Engineering, The University of Western Ontario, London, Ontario, Canada, N6A 3K7; Department of Medical Biophysics, Schulich School of Medicine and Dentistry, The University of Western Ontario, London, Ontario, Canada, N6A 5C1; Robarts Research Institute, The University of Western Ontario, London, Ontario, Canada, N6A 5C1; Department of Chemical Engineering, Faculty of Engineering, Queen’s University, Kingston, Ontario, Canada, K7L 3N6; Department of Physiology and Pharmacology, Schulich School of Medicine and Dentistry, The University of Western Ontario, London, Ontario, Canada, N6A 5C1; Department of Chemical & Biochemical Engineering, Faculty of Engineering, The University of Western Ontario, London, Ontario, Canada, N6A 5B9; Department of Anatomy and Cell Biology, Schulich School of Medicine and Dentistry, The University of Western Ontario, London, Ontario, Canada, N6A 5C1

**Keywords:** Cell therapy, vascular regeneration, adipose-derived stromal cells (ASC), hydrogels, chitosan, host response

## Abstract

The delivery of human adipose-derived stromal cells (hASCs) to ischemic tissues represents a promising strategy to promote vascular regeneration for patients with critical limb ischemia (CLI). Building on previous work, this study focused on the *in vivo* characterization of a hydrogel cell delivery platform for hASCs composed of peptide-functionalized methacrylated glycol chitosan (MGC-RGD) and a terminally acrylated triblock copolymer of poly(ethylene glycol) and poly(trimethylene carbonate) (PEG(PTMC-A)_2_) in athymic *nu/nu* mice with femoral artery ligation-induced critical limb ischemia (FAL-CLI). This immunodeficient mouse strain was selected to enable human cell transplantation in a model with conserved monocyte/macrophage function, recognizing that macrophages are key regulators of the biomaterial implant response, as well as vascular repair and regeneration. The hASCs were engineered to co-express tdTomato and firefly luciferase to enable longitudinal cell tracking using bioluminescence imaging (BLI). Interestingly, the hASCs were better retained following delivery in saline compared to hydrogel delivery. However, laser Doppler perfusion imaging (LDPI) analysis indicated that the restoration of hindlimb perfusion was similar between the two cell treatment groups. Critically, delivery of the hASCs within the hydrogels was associated with adverse outcomes only observed within this treatment group, including severe swelling, discoloration, and necrosis, which necessitated early euthanasia of some mice. CD45 staining supported that the combination of the cells and the hydrogels induced an inflammatory host response. These findings contrast with previous positive results when the platform was tested for hASC delivery in more severely immunocompromised NOD/SCID mice with FAL-CLI, as well as allogeneic rat ASC delivery in a healthy immunocompetent rat model. Overall, this study emphasizes the potential importance of testing cell delivery platforms in pre-clinical disease models that have retained host immune cell function, especially for immunomodulatory cell populations such as ASCs.

## 1 Introduction

Atherosclerotic cardiovascular diseases remain the leading cause of mortality worldwide, with peripheral arterial disease (PAD) being the third leading cause of atherosclerotic morbidity, afflicting more than 230 million people worldwide [1], [2]. Critical limb ischemia (CLI) represents the end stages of PAD and is associated with symptoms such as limb pain at rest, non-healing ulcers, and gangrene [3]. Cell therapies for PAD involving the localized delivery of mesenchymal stromal cells (MSCs) have attracted significant interest due to their capacity to stimulate angiogenesis, modulate inflammation and promote tissue regeneration through paracrine-mediated mechanisms [4]. In particular, adipose-derived stromal cells (ASCs) represent a promising clinically-translational cell population that is more accessible and abundant than other MSC sources [5]. Notably, recent evidence suggests that ASCs may have higher pro-angiogenic regenerative potential than MSCs derived from bone marrow [6], [7].

Although delivery of MSCs has shown success in preclinical models of CLI, clinical trials to date have not clearly demonstrated the efficacy of MSC-based therapies for this application [8]. Postulated reasons for this gap in clinical translation include the poor survival and retention of the cells following directed delivery to the harsh environment within ischemic muscle tissue [9]–[11]. Seeking to address this barrier, there has been growing interest in the application of hydrogels as cell delivery platforms for vascular regeneration, including the treatment of PAD [12], [13]. Previous work from our lab developed a mechanically-resilient composite hydrogel delivery system to support the survival and pro-angiogenic capacity of ASCs as a therapy for CLI [11], [14]. Specifically, this hydrogel was composed of crosslinkable methacrylated glycol chitosan functionalized with the fibronectin-derived cell adhesive peptide motif arginine-glycine-aspartate (MGC-RGD), and a terminally acrylated triblock copolymer of poly(ethylene glycol) and poly(trimethylene carbonate) (PEG(PTMC-A)_2_) [15]. The MGC-RGD was chosen to provide a cell-supportive environment, while the PEG(PTMC-A)_2_ was incorporated to match the mechanical requirements of muscle tissue by providing a structural component capable of sustaining large, repeated mechanical deformations [11], [15]. Human ASCs (hASCs) showed enhanced retention at 28 days when injected intramuscularly within the hydrogels to a femoral artery ligation-induced CLI (FAL-CLI) model in immunodeficient NOD/SCID mice as compared to delivery in saline, based on end-point analyses of human leukocyte antigen-ABC^+^ (HLA-ABC^+^) cells in tissue cross-sections [11]. Further, hydrogel delivery significantly enhanced the endothelial cell density in the ischemic gastrocnemius muscles compared to hASCs delivered in saline, as well as with hydrogel and saline alone controls [11].

The previous study was performed using NOD/SCID mice, selected based on their capacity to support human cell engraftment and function without xenorejection [16]. Acknowledging the importance of the host immune response in mediating tissue regeneration, follow-up testing was performed in a healthy immunocompetent rat model [14]. Interestingly, intramuscular (IM) delivery of allogeneic rat ASCs within the hydrogels augmented the recruitment and infiltration of host CD68^+^ macrophages relative to hydrogel alone controls. Moreover, significantly more CD31^+^ vessels were observed within and surrounding the hydrogels that incorporated rat ASCs at both 2 and 4 weeks following IM injection. Taken together, these results demonstrated that ASCs could modulate the host macrophage response to the hydrogels, and that this may be favorable for augmenting vascular regeneration.

Building from our previous work, this study sought to perform additional characterization of the MGC-RGD+PEG(PTMC-A)_2_ delivery system using clinically-relevant human ASCs within the FAL-CLI model in athymic *nu/nu* mice, which have T-lymphocyte deficiency with normal monocyte/macrophage content, in contrast to the NOD/SCID mice that are known to have deficiencies in macrophage number and function [16], [17]. Recognizing the value of non-invasive imaging technologies to quantitatively assess vascular regeneration and to simultaneously track the survival of therapeutic cell populations, laser Doppler perfusion imaging (LDPI) was integrated to assess changes in hindlimb perfusion over time, along with bioluminescence imaging (BLI) to longitudinally track the hASCs that were engineered through lentivirus transduction to stably co-express firefly luciferase (FLuc) and tdTomato (tdT). Based on our previous work, it was hypothesized that delivery within the hydrogels would augment the localized retention of viable hASCs as compared to delivery in saline in the *nu/nu* mice, and that this would be coupled with accelerated restoration of hindlimb perfusion, indicative of enhanced vascular regeneration.

## 2 Results

### 2.1 In Vitro Cell Encapsulation

hASCs were encapsulated using a thermally-induced crosslinking approach and cultured for up to 7 days *in vitro* within the composite hydrogels. As shown in Figure 1, the encapsulated hASCs remained viable across the 7-day culture period, with their morphology showing increased spreading over time, consistent with our previous work [11].

**Figure 1.**
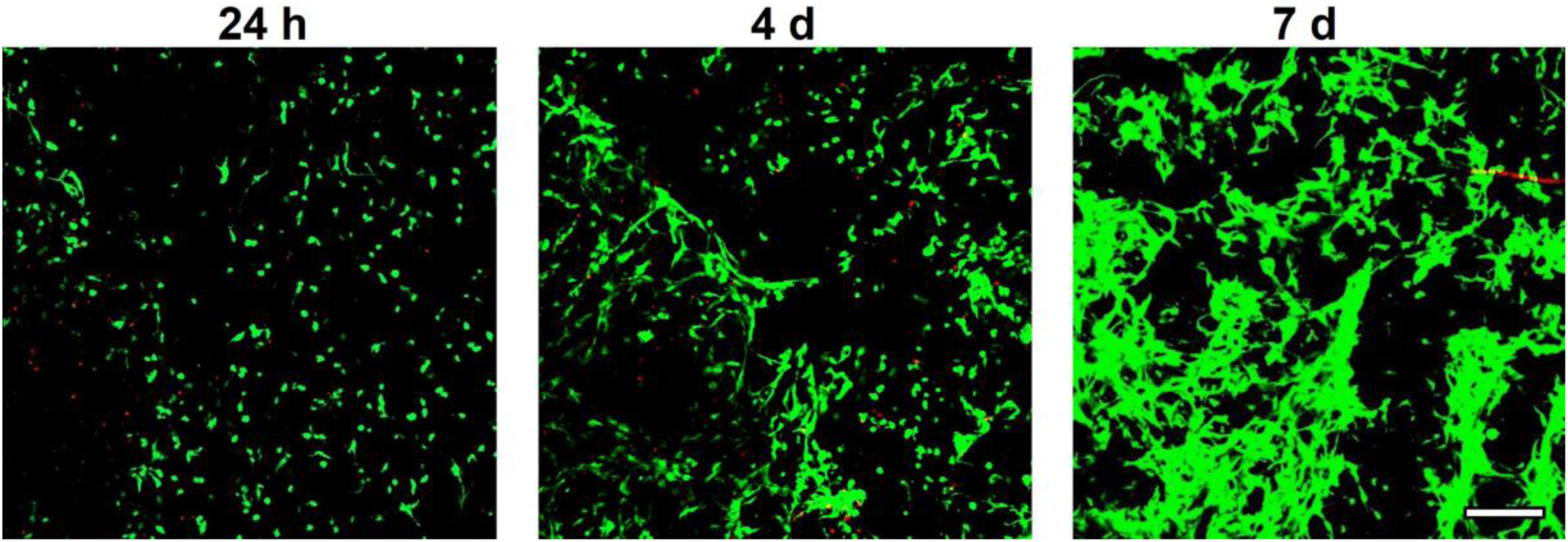
hASCs encapsulated in MGC-RGD+PEG(PTMC-A)_2_ hydrogels remained viable over 7 days of culture, showing increased spreading over time. Live/Dead™ staining was used to visualize hASCs within the composite hydrogels. Representative confocal microscopy imaging shows viable calcein^+^ cells in green, and dead EthD1^+^ cells in red. Scale bar represents 500 μm.

To enable longitudinal *in vivo* cell tracking via BLI, hASCs were engineered through lentiviral transduction to stably co-express FLuc to facilitate detection via BLI and tdT to identify the cells via flow cytometry and in tissue cross-sections [18]. Flow cytometry confirmed a high transduction efficiency (>85%) (Supplementary Figure S1). In addition, flow cytometry showed that transduction did not alter cell viability (>95%) or the hASC immunophenotype (Supplementary Figure S1). *In vitro* studies using IVIS imaging to quantitatively assess luminescence associated with the transduced hASCs confirmed the sustained presence of viable hASCs following encapsulation within the composite MGC-RGD+PEG(PTMC-A)_2_ hydrogels and culture for up to 14 days, with no significant difference in the signal over time (Figure 2).

**Figure 2.**
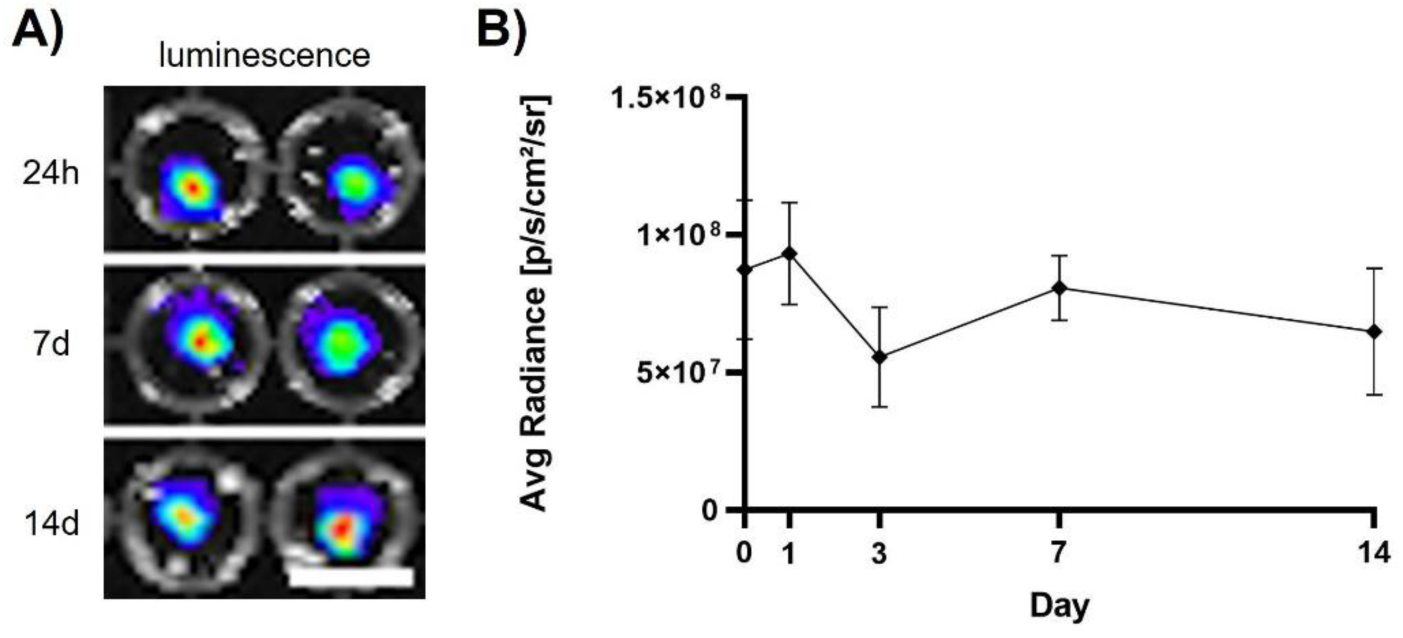
*In vitro* luminescence imaging confirmed high hASC viability following hydrogel encapsulation and culture over 14 days. (A) Representative images show luminescent signal from the transduced FLuc^+^tdT^+^ hASCs encapsulated and cultured in the MGC+PEG(PTMC-A)_2_ hydrogels. Scale bar = 5 mm. (B) Luminescent signals quantified across the 14-d culture period. All data presented as mean ± SD. No significant differences in average radiance across timepoints were detected using one-way ANOVA with Tukey’s post-hoc test (n=4 hydrogels/timepoint).

### 2.2 Characterization of hASC delivery in the MGC-RGD + PEG(PTMC-A)_2_ hydrogels in the FAL-CLI model in athymic nude (nu/nu) mice

The FAL-CLI model was applied in athymic nude mice to compare the efficacy of delivering the transduced hASCs in the composite MGC-RGD+PEG(PTMC-A)_2_ hydrogels to hASC delivery in saline, with hydrogel alone and saline alone groups included as controls. Treatments were delivered at 24 h post-ligation, with all mice receiving 4x10^5^ hASCs (2x10^7^ cells/mL) in 20 μL injections. Mice were assessed over 35 days, with regular BLI to assess hASC viability and retention and LDPI to assess recovery of perfusion. Testing was completed on a total of 3 cohorts, with 6 mice per cohort, and each cohort was treated with a different hASC donor. Only the data from the first two cohorts are included in Figures 3 and 4, as the mice treated with the hASCs in the MGC-RGD+PEG(PTMC-A)_2_ hydrogels in the third cohort had to be euthanized early due to adverse events, further detailed below.

**Figure 3.**
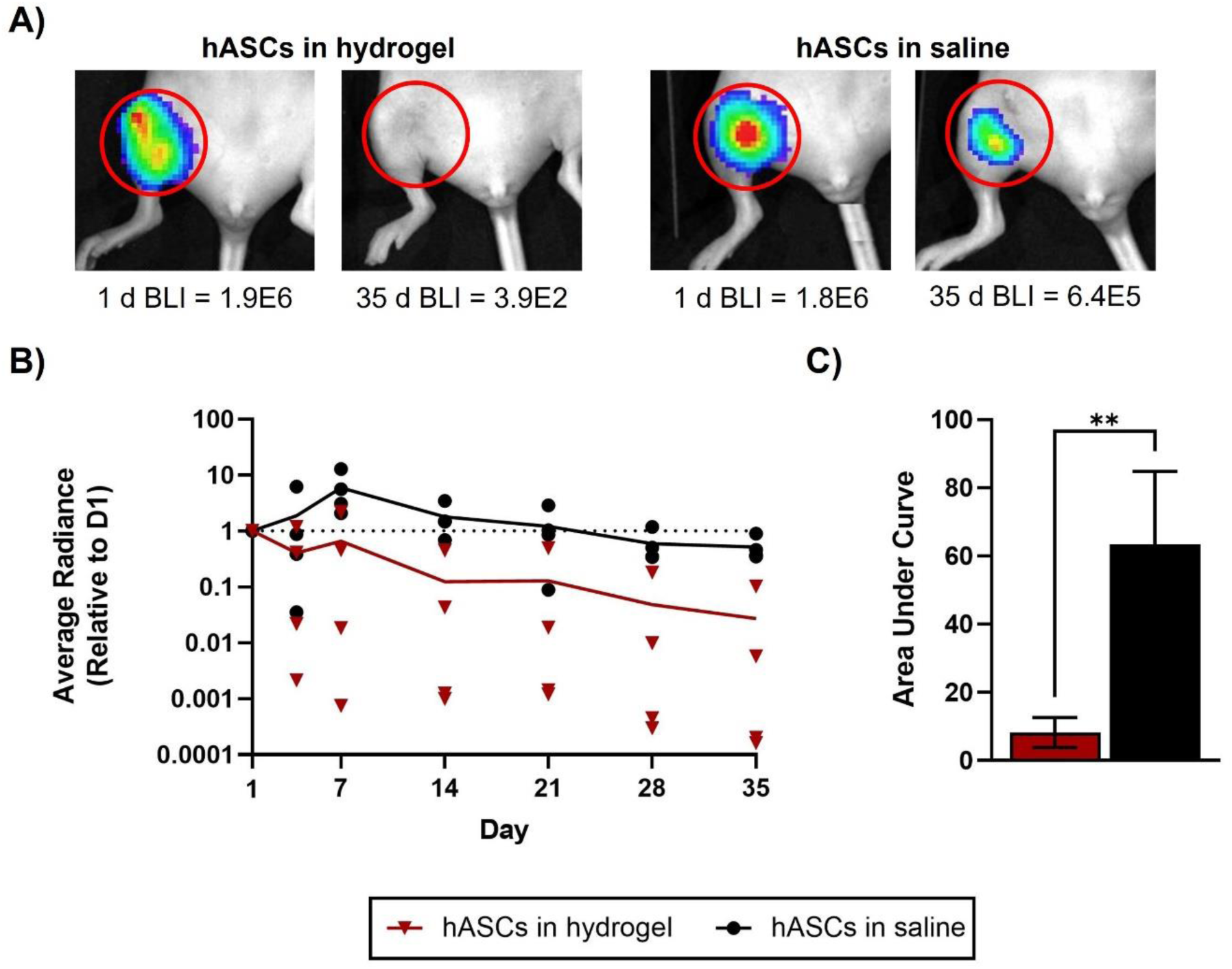
hASCs were better retained in the FAL-CLI model in *nu/nu* mice when delivered in saline as compared to the MGC-RGD+PEG(PTMC-A)_2_ hydrogels. (A) Representative BLI images showing FLuc^+^ hASC signal in mice that received hASCs delivered in the hydrogel or saline at day 1 and 35. Each mouse received 100 µL IP injection of D-luciferin, and bioluminescent images were taken until the signal peaked within the region of interest (ROI). (B) Average bioluminescent radiance (p/s/cm^2^/sr) was calculated within an ROI around the injection site for each mouse at each timepoint, and normalized to day 1. Individual data points for each mouse are shown, with the group mean plotted as a solid line. Dotted line represents day 1 values. Normalized average radiance for mice that received hASCs in hydrogel showed variability; one mouse had ∼10% retention at day 35, and one mouse had ∼1% retention, while two mice showed negligible retained cells. No statistically significant differences were detected between treatment groups at any timepoint, using multiple unpaired t-tests with a false discovery rate set to 5% (n=4 mice/treatment group). (C) AUC analysis of the normalized BLI data, indicating that the hASCs delivered in saline were better retained than the hASCs delivered using the hydrogel. Data presented as mean ± SD. Differences in the AUC were assessed using an unpaired t-test (n=4 mice/treatment group); Significant differences are indicated (** p<0.01).

**Figure 4.**
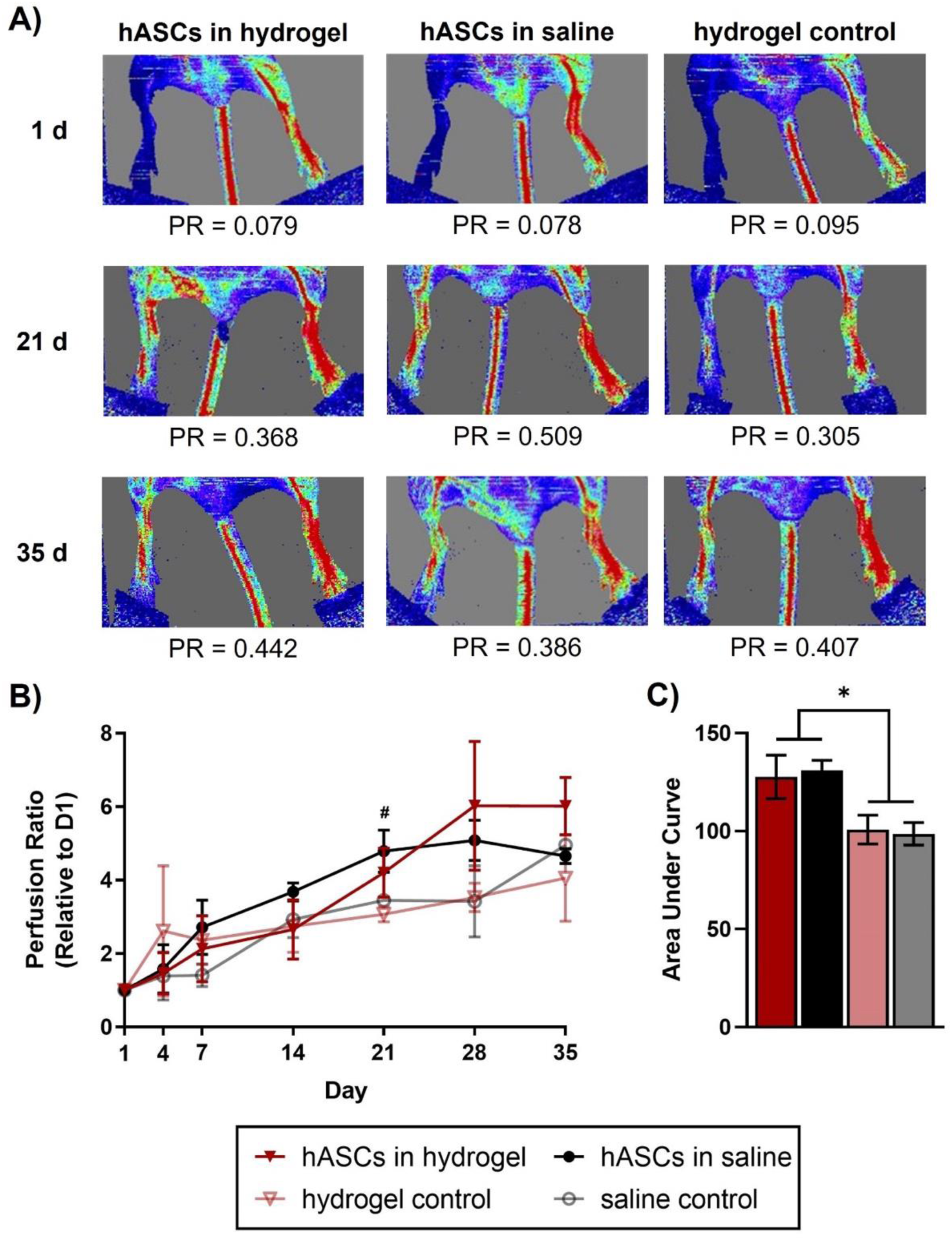
Delivery of hASCs in *nu/nu* mice with FAL increased the recovery of limb perfusion. (A) Representative LDPI images showing blood flow in the surgical and control limbs at days 1, 21, and 35 after the delivery of hASCs in hydrogel or saline, or hydrogel controls. (B) Mean flux was quantified within an ROI comprising the foot and ankle for both limbs. Perfusion ratio was calculated from the mean flux of surgical limb / control limb for each mouse at each time point, and normalized to the perfusion ratio at day 1. At day 21, mice that received hASCs in saline had significantly higher perfusion compared to mice that received the hydrogel or saline alone controls (# p<0.05). This data encompasses two sets of surgeries taken out to 35 days (n=4 mice/hASC treatment group, n=2 mice/control treatment group). Data presented as mean ± SD. Differences between treatment groups at each timepoint were detected using two-way RM-ANOVA, with Tukey’s post-hoc test for multiple comparisons. (C) Area under the curve (AUC) was calculated from the perfusion ratio data. Both hASC treatment groups resulted in significantly higher AUC compared to both control groups (* p<0.05), indicating that hASC treatment accelerated the recovery of limb perfusion in the FAL-CLI model in nu/nu mice. Data presented as mean ± SD. Differences between treatment groups were detected using one-way ANOVA, with Tukey’s post-hoc test for multiple comparisons (n=4 mice/hASC treatment group, n=2 mice/control treatment group).

Interestingly, the BLI data suggested that hASCs delivered in saline were better retained in the *nu/nu* model compared to hASCs delivered in the hydrogel (Figure 3). Of note, BLI measurements for mice that received the hASCs in saline appeared to increase from day 1 to 7, which may be attributed to enhanced delivery of the D-luciferin to the ischemic region. All mice that received hASCs delivered in saline showed >35% retention at day 35 compared to day 1, while all mice that received hASCs delivered in hydrogel showed ≤10% retention at day 35 compared to day 1, with one mouse showing 10% retention, and the other three showing ≤0.6% retention. Area under the curve (AUC) analysis showed a significant difference between the groups, supporting that cell retention was enhanced when the hASCs were delivered in saline compared to delivery within the MGC-RGD+PEG(PTMC-A)_2_ hydrogels (Figure 3C).

Complementary LDPI indicated that the mice that received hASCs delivered in saline had significantly increased hindlimb perfusion at day 21 compared to the hydrogel alone and saline alone controls (Figure 4B). However, AUC analysis of the LDPI data plots over 35 days suggested that hASC delivery, either within the hydrogels or saline, significantly increased the overall recovery of limb perfusion compared to the hydrogel alone and saline alone controls (Figure 4C).

Notably, several adverse events were observed in mice that were treated with the hASCs in the MGC-RGD+PEG(PTMC-A)_2_ hydrogels, which weren’t observed in any other treatment group. Table 1 summarizes the specific adverse events and the timepoints at which they were observed. Interestingly, mice that received the hASCs within MGC-RGD+PEG(PTMC-A)_2_ hydrogels frequently demonstrated discoloration, swelling, and possible capsule or cyst formation at the site of injection, along with stiffness, difficulty ambulating, and discoloration of the toes or toenails. These symptoms were not observed in the hydrogel alone controls, or when the cells were delivered in saline, indicating that it was the combination of the cells and the hydrogel that was causing the adverse events. In the third cohort, toe necrosis and early signs of auto-amputation were observed in the two mice that received hASCs within the MGC-RGD+PEG(PTMC-A)_2_ hydrogels, which necessitated early euthanasia at day 12.

**Table 1.** Summary of adverse events for mice that received hASCs in hydrogel treatment.

| Symptom | Timepoint |  |  |  |  |
| --- | --- | --- | --- | --- | --- |
|  | D4 | D7 | D12/<br>D14 | D21 | D28 |
| **Discoloration @ injection site (dark purple)** | 4/6 | 4/6 | 5/6 | 2/4 | 2/4 |
| **Discoloration of toes / toenails** | 1/6 | 2/6 | 3/6 | 0/4 | 0/4 |
| **Necrosis of toes, signs of auto-amputation** | 0/6 | 0/6 | 2/6 | 0/4 | 0/4 |
| **Cyst / Capsule @ injection site** | 0/6 | 2/6 | 4/6 | 2/4 | 2/4 |
| **Edema** | 4/6 | 5/6 | 4/6 | 2/4 | 2/4 |
| **Stiffness** | 0/6 | 0/6 | 2/6 | 2/4 | 2/4 |
| **Difficulty ambulating** | 1/6 | 1/6 | 4/6 | 2/4 | 2/4 |
| **Early euthanasia required** | 0/6 | 0/6 | 2/6 | 0/4 | 0/4 |

Immunofluorescence (IF) staining was performed for murine CD45 in thigh muscle sections of the two cohorts of mice that were euthanized at day 35, as well as the cohort euthanized at the earlier timepoint (Figure 5A & Supplementary Figure S2). Semi-quantitative scoring was performed in a blinded manner based on the following scale: 0 (no dense CD45^+^ staining outside of the ligated vessel region), 1 (only one small region of dense CD45^+^ staining), 2 (one medium region of dense CD45^+^ staining or multiple small regions) or 3 (at least one large region of dense CD45^+^ staining). Although there were only a small number of mice analyzed at the early timepoint, regions of dense CD45^+^ staining were only observed in the mice that received hASCs delivered in the hydrogel (Figure 5B), consistent with the discoloration, edema, and swelling in these mice that necessitated euthanasia. Overall, the results at both timepoints indicated that hASC delivery using the hydrogel was more frequently associated with a more potent inflammatory response.

**Figure 5.**
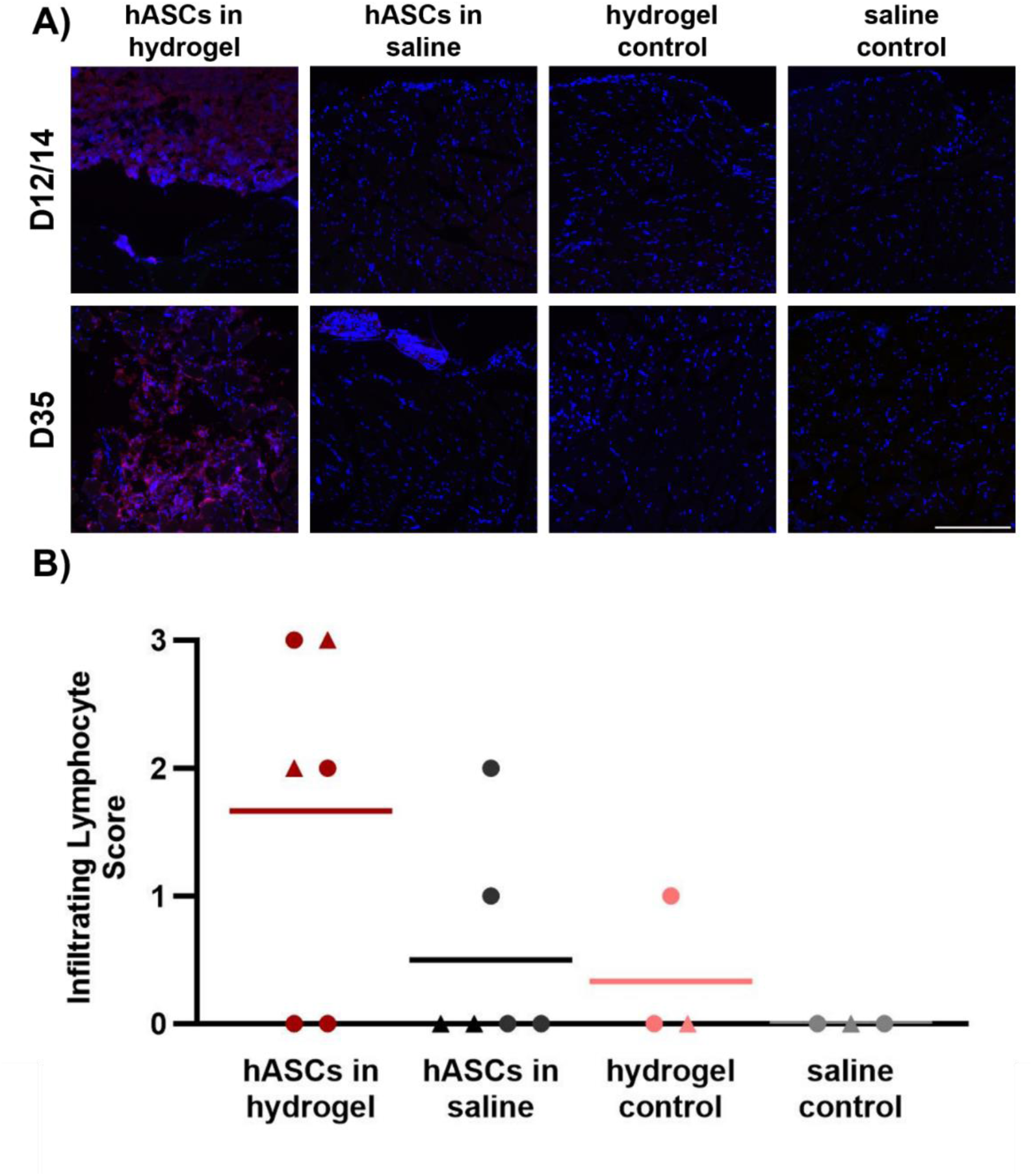
CD45^+^ staining showed a more potent inflammatory response in a higher fraction of the mice within the muscles treated with the hASCs in the hydrogel. Large regions containing a dense population of CD45^+^ cells were only observed in mice that received hASCs delivered in the hydrogel. (A) Representative IHC images of mouse thighs harvested post-euthanasia at days 12/14 or 35 following FAL (DAPI^+^ cells in blue and CD45^+^ cells in red). Scale bar represent 250 μm. Larger tiled images used for scoring are shown in Supplementary Figure S2. (B) Semi-quantitative assessment of regions of dense CD45^+^ staining within the thigh muscles of the ligated limb was performed, with blinded scoring on a scale of 0-3. Graph includes pooled data of all samples collected post-euthanasia at days 12/14 (triangles) and 35 (circles). Data points represent scoring for individual mice, with the line representing the mean score for each treatment group.

## 3 Discussion

Immunocompromised mouse models are an important tool in the preclinical evaluation of human cell-based therapies [26]. In particular, NOD/SCID mice are frequently used in studies where long-term cell engraftment is desired, due to complete ablation of T- and B-lymphocytes (SCID mutation), and reduced macrophage numbers and function (NOD mutation) [19]. However, endogenous host immune cells are important mediators of both inflammation and vascular regeneration [20]. In particular, monocytes and macrophages play key roles in mediating angiogenesis and inflammation [21], [22]. Moreover, it has been suggested that the full therapeutic effects of MSCs require the presence of endogenous macrophages/monocytes [23]. For this reason, the primary focus of the current study was to characterize the response of hASCs delivered within the MGC-RGD+PEG(PTMC-A)_2_ hydrogels in the FAL-CLI model in athymic *nu/nu* mice. This model was selected because while *nu/nu* mice lack a T-cell response, they have more functional macrophages compared to the more severely immunocompromised NOD/SCID model. Interestingly, when the FAL-CLI model was applied in athymic *nu/nu* mice, AUC analysis for the LDPI showed a significant increase for mice that were treated with hASCs, independent of the delivery method, compared to the hydrogel alone and saline alone controls. These findings contrast with our previous histological data in the NOD/SCID model [11], which suggested that vascular regeneration was augmented when the hASCs were delivered within the hydrogels.

Recognizing that methods to track the survival, retention, and fate of delivered cells are essential in the development of MSC-based cell therapies to better understand the underlying mechanisms of regeneration, we engineered the hASCs to express firefly luciferase to enable longitudinal *in vivo* cell tracking by BLI [24]. BLI offers notable advantages over the more standard practice of using cell-specific markers to detect the delivered cell populations within tissue cross-sections, which is extremely time-consuming and challenging to accurately quantify, and can only be performed at end-point after the cells may have already been cleared [24]. Importantly, only viable cells will produce luciferase and the luciferase-luciferin reaction requires ATP to produce light, and thus BLI detects live cells and can be used to track the survival and location of transplanted hASCs within individual mice over time [25]. Interestingly, the BLI data indicated that hASC retention/survival was augmented when the cells were delivered in saline as compared to within the hydrogels. Again, these findings are in contrast to the previous study in the NOD/SCID mice, where the hASCs were detected based on staining for HLA-ABC^+^ cells in the muscle at 4 weeks post-transplantation when delivered in the hydrogels, but not when delivered in saline [11]. Although the variations in the cell tracking methods used in the two studies prevent direct comparison between the two mouse strains, the findings suggest that differences in the host response to the hASCs delivered in the MGC-RGD+PEG(PTMC-A)_2_ hydrogels impaired the retention of viable hASCs within the *nu/nu* mice.

Compared to studies in the literature that have indicated there is low cell retention following delivery in saline, the BLI data indicated that >35% of the hASCs delivered in saline were retained within the thigh muscles at the 35-day timepoint in the athymic *nu/nu* mice. It is important to note that differences across studies could be related in part to the specific MSC source applied, including whether the delivered cells are syngeneic, allogeneic, or xenogeneic in nature and if immunocompetent or immunocompromised mice are used. Unfortunately, comparative data is limited as only a small number of studies to date have characterized cell retention when developing cell therapies for CLI. In contrast to our findings, Laurila *et al*. delivered 1x10^6^ Luc^+^ MSCs derived from human embryonic stem cells in saline to an FAL-CLI model in immunocompetent Fischer 344 rats and showed using BLI that the relative luminescent signal decreased to 29% at 6 h post-delivery, and further decreased to only 1.5% of the original intensity by 24 h post-delivery [10]. Using longitudinal fluorescent imaging, Wang *et al.* showed that the signal intensity of mouse ASCs delivered in saline to an FAL-CLI model in BALB/c nude mice decreased gradually to background levels by day 14 [26]. Also applying the FAL-CLI model in BALB/c nude mice, Fan *et al*. used BLI to demonstrate ∼25% of FLuc^+^ mouse ASCs were retained on day 14 when delivered in saline, followed by a progressive decline to background levels over 6 weeks [27]. Interestingly, Fan *et al*. also used longitudinal fluorescent imaging to track the cells, which showed a more rapid decrease in signal to background levels by day 28, confirming that different imaging modalities can influence the detection of retained cells [27].

While BLI is a powerful approach for tracking cell retention and survival, it is important to note that the impaired delivery of the D-luciferin is a potential limitation of its use in the FAL-CLI model, especially during the early stages following surgery when blood flow to the ischemic limb remains highly restricted. It would be expected that as vascular regeneration progresses and hindlimb perfusion is restored, there would be augmented and/or accelerated delivery of the luciferin substrate, which could impact the signal detected, as was observed with the relative increase in signal from day 1 to 7 in the hASC in saline group in the current study. While this makes absolute quantification of cell retention challenging at early time points, BLI remains a useful tool for comparing hASC retention between the two different delivery methods applied in the current study at each respective timepoint. As an alternative substrate-free approach, future studies could explore the use of near-infrared fluorescent reporters, which would allow for improved depth of penetration of signal compared to other fluorescent reporters and have shown promise for longitudinal *in vivo* tracking of delivered cells [28].

Combining the LDPI and BLI data, our findings suggest that enhanced cell retention may be insufficient to augment vascular regeneration. These results are in contrast to our previous findings in the NOD/SCID mice [11], as well as the previously-mentioned work of Wang *et al*., who showed that delivery of mouse ASCs within composite chitosan-hyaluronic acid-based hydrogels enhanced their retention compared to delivery in saline, and that this was correlated with the increased recovery of hindlimb perfusion based on LDPI [26]. Notably, when the hASCs were delivered in the MGC-RGD+PEG(PTMC-A)_2_ hydrogels in the FAL-CLI model in athymic *nu/nu* mice, adverse events consistent with a negative host inflammatory response were observed in a higher fraction of the mice, with 5/6 mice showing marked swelling and limb discoloration, and 2/6 having a severe response including tissue necrosis that necessitated euthanasia. Importantly, none of these adverse events were observed in any of the other treatment groups, indicating that it was the combination of the cells and the hydrogel that was contributing to the negative immunological response.

Our conflicting findings, as well as previous work from other groups, highlight the influence of animal model selection on the apparent efficacy of cell-based therapies. For example, Brady *et al*. showed differences in cell survival and blood vessel formation following implantation of engineered tissues seeded with human pluripotent stem cell-derived cardiomyocytes in athymic nude mice versus athymic nude rats [29]. Importantly, mouse strain has been shown to influence the severity of ischemia [30], [31], which may have also contributed to a harsher and more inflammatory microenvironment in the athymic mouse model used in the current study as compared to the previous NOD/SCID model. In particular, a high incidence of ischemic limb necrosis has been reported in nude mice, which although not yet well understood, is thought to be related to the immune response and the ability to form collaterals [30], [31]. Based on the adverse outcomes encountered, further testing of our platform within this model was discontinued for ethical reasons.

Consistent with the macroscopic signs of a negative host inflammatory response, IF staining revealed dense regions of infiltrating CD45^+^ murine cells within the ischemic muscles in a higher fraction of the mice that received the hASCs delivered in the hydrogel. We hypothesize that the hASCs within the hydrogels stimulated a potent host macrophage response in the *nu/nu* mice, which may have led to accelerated hASC clearance. This interpretation is supported by our previous studies of the MGC-RGD+PEG(PTMC-A)_2_ hydrogels in the healthy immunocompetent rat model, which showed enhanced recruitment and greater infiltration of CD68^+^ macrophages when rat ASCs were delivered within the hydrogels as compared to hydrogel alone controls, with most of the macrophages expressing the pro-inflammatory “M1-like” marker CCR7 [14]. This macrophage response was associated with enhanced blood vessel formation rather than adverse outcomes, which could be related at least in part to differences in the microenvironment within healthy muscle versus the muscles affected by the FAL surgery. Similarly, in addition to rapid MSC clearance from the site of injection, Laurila *et al*. showed increased CD68^+^ macrophage infiltration in rats that received MSCs compared to saline controls in the FAL-CLI model in immunocompetent rats, suggesting that the macrophages may have contributed to the low cell survival or retention observed in their study [10].

Moving forward, the application of humanized mouse models is warranted to better assess how delivered human MSCs will interact with human immune cells and modulate the inflammatory response, including systematically probing the effects of different delivery strategies and cell doses [32], [33]. In addition, it would be interesting to investigate clodronate-induced macrophage depletion in mouse models to more specifically assess the roles of macrophages in mediating the response to cell-based therapies for vascular regeneration [34].

## 4 Conclusion

Overall, the results of this study in comparison to our previous characterization of the MGC-RGD+PEG(PTMC-A)_2_ hydrogels highlight how differences in the host immune system and/or the induction of ischemia in rodent models can profoundly influence the *in vivo* efficacy of biomaterials-based cell delivery for vascular regeneration. In contrast to our previous studies comparing ASC delivery intramuscularly in the hydrogels or saline in NOD/SCID mice with FAL-CLI or healthy immunocompetent rats, the hASCs delivered intramuscularly to athymic *nu/nu* mice with FAL-CLI were better retained following delivery in saline as compared to hydrogel delivery. Moreover, the combined LDPI and BLI data indicated that enhancing hASC retention was not sufficient to augment vascular regeneration. Notably, a negative immune response was observed when the hASCs were delivered within the hydrogels in the athymic *nu/nu* mice, which were not observed in the other treatment groups. Overall, these findings suggest that the MGC-RGD+PEG(PTMC-A)_2_ hydrogels should not be used as a delivery vehicle for hASCs in a highly pro-inflammatory environment. When assessing MSC delivery using biomaterials, the interactions between the delivered cells, the materials, and the host cells must be considered and carefully evaluated. As biomaterials can influence the host immune response, as well as the function of delivered cells, it will be essential to perform testing of human cell therapies in relevant disease models that apply animals with functional immune systems to better recapitulate the human response and develop strategies that have a higher likelihood of success when translated to the clinic.

## 5 Methods

### 5.1 Materials

Unless otherwise stated, all chemicals and reagents were purchased from Millipore Sigma, Canada, and used as received.

### 5.2 Polymer synthesis

Polymers composed of MGC-RGD and PEG(PTMC-A)_2_ were synthesized according to previously published protocols [11], [15]. For PEG(PTMC-A)_2_ synthesis, PEG(PTMC)_2_ was first prepared through the ring-opening polymerization of trimethylene carbonate (TMC) (LEAPChem), initiated by PEG diol (PEG_20_, M_N_ = 20 kDa) . The molecular weight of the polymer was assessed using ^1^H NMR in deuterated DMSO (DMSO-d_6_) on an Inova 600 NMR spectrometer (Varian), confirming the molar ratios of PEG_20_(PTMC_2_)_2_. PEG(PTMC)_2_ was subsequently acrylated using acryloyl chloride to a high degree of acrylation (85%), consistent with our previous work [11], as confirmed by ^1^H NMR spectroscopy (Supplementary Figure S3A).

MGC was synthesized through the methacrylation of glycol chitosan (GC) (minimum M_N_ = 82 kDa, 85% degree of deacetylation, Wako Chemicals Inc.) using glycidyl methacrylate, followed by ^1^H NMR spectroscopy in deuterium oxide (D_2_O) at 80°C to assess the degree of methacrylation. For peptide conjugation to generate the MGC-RGD, GGGRGDS peptide (>90% purity, CanPeptides Inc.) was first acrylated using N-acryloxysuccinimide, with ^1^H NMR in D_2_O at 80°C used to assess the degree of acrylation. Subsequently, the acrylated RGD peptide was reacted with the MGC polymer to obtain MGC-RGD with a degree of substitution of 5%, consistent with our previous work [11], as confirmed by ^1^H NMR spectroscopy (Supplementary Figure S3B).

### 5.3 Isolation and culture of hASCs

Resected human adipose tissue was obtained with informed consent from routine breast or abdominal reduction surgeries conducted at the London Health Sciences Centre in London, ON, Canada, with approval from the Human Research Ethics Board at Western University (HSREB# 105426). Fresh adipose tissue samples were transported to the lab on ice in sterile phosphate buffered saline (PBS) (Wisent Inc) supplemented with 2% bovine serum albumin (BSA) (Bio-shop) and hASCs were isolated according to published methods and cultured in ASC proliferation medium consisting of DMEM/F12 supplemented with 10% FBS and 1% penicillin-streptomycin (Gibco™) [35]. Media was replaced every 2-3 days, and passaging was performed at 80% confluence. hASCs were frozen at passage 0 and stored in liquid nitrogen until needed, with hASCs at passage 3-5 used for all studies. Cell donor information is summarized in Supplementary Table S1.

### 5.4 hASC transduction

To enable longitudinal *in vivo* cell tracking, passage 1 hASCs were engineered through lentiviral transduction to co-express tdT and codon-optimized firefly luciferase (FLuc2) under the human elongation factor 1-α promoter for constitutive expression (pEF1-alpha-tdT-FLuc2 reporter), according to previously published protocols, with a viral multiplicity of infection (MOI) of 75 and 100 μg/mL protamine sulphate [18].

### 5.5 Assessment of immunophenotype of transduced hASCs

Flow cytometry analyses were performed to assess the viability, immunophenotype, and tdT expression of the transduced hASCs in comparison to non-transduced controls. Passage 2 hASCs were extracted using Versene solution (Gibco™), with a 10 min incubation on ice, followed by a 10 min incubation at 37°C. Remaining adherent cells were collected in PBS using a cell scraper, and hASCs were filtered through a 100 μm filter prior to downstream staining. Samples were stained and assessed for expression of CD90, CD29, CD146, CD31 and CD45 as outlined in Table 2. Cell viability was assessed using the Live/Dead™ Fixable Aqua kit (Invitrogen™), with a 1:500 dilution. tdTomato signal was compared to non-transduced cells. Samples were analyzed with an LSR II Flow Cytometer (BD Biosciences) at the London Regional Flow Cytometry Facility. Data was analyzed using FlowJo v10 with gates set using the relevant fluorescence minus one (FMO) controls.

**Table 2.** List of antibody dilutions for hASC flow cytometry characterization.

| Antigen | Conjugation | Dilution | Catalog Number | Company |
| --- | --- | --- | --- | --- |
| CD90 | BV605 | 1:800 | 328128 | Biolegend |
| CD29 | AF488 | 1:100 | 303016 | Biolegend |
| CD146 | APC-Fire™750 | 1:50 | 361028 | Biolegend |
| CD31 | PE-Cy7 | 1:400 | 303118 | Biolegend |
| CD45 | APC | 1:200 | 368532 | Biolegend |

### 5.6 hASC encapsulation for in vitro studies

Hydrogels were prepared according to previously described protocols and formulations [11]. In brief, dry MGC-RGD and PEG(PTMC-A)_2_ were decontaminated under low-intensity UV light in a biosafety cabinet for 30 min. The weight of each polymer was calculated based on a final formulation of 1% (w/v) MGC-RGD and 4% (w/v) PEG(PTMC-A)_2_. The polymers were dissolved in sterile PBS (75% of the final volume) overnight at room temperature under constant agitation on an orbital shaker at 100 RPM.

hASCs were extracted using trypsin (Wisent Inc.), resuspended at 5x10^7^ cells/mL (5X final concentration) in ASC proliferation medium (20% of the final volume) and thoroughly combined with the pre-polymer solution. Concentrated solutions of ammonium persulphate (APS) (Bio-Shop) and tetramethylethylenediamine (TEMED) (Bio-Shop) were sterile filtered and added sequentially to the combined mixture with thorough mixing with the pipette tip between each addition, to a final concentration of 5 mM each of APS and TEMED. The volumetric ratio of the final solution was 75% pre-polymer solution : 20% hASC suspension : 2.5% APS : 2.5% TEMED, with a final cell density of 10^6^ cells/mL. Hydrogels for the *in vitro* cell culture studies were prepared in syringe molds using 0.3 mL insulin syringes, and allowed to gel at 37°C for 15 min. Following gelation, 10 μL hydrogels were transferred to individual wells of a 12-well plate, and cultured in 2 mL of ASC proliferation medium, with media changes every 2 days.

### 5.7 In vitro assessment of cell viability

Live/Dead™ staining was performed at 24 h, 4 d, and 7 d post-encapsulation to assess the viability of non-transduced hASCs encapsulated within the MGC-RGD+PEG(PTMC-A)_2_ hydrogels. Hydrogels were first rinsed with PBS, and then incubated in a solution of 2 μM Calcein-AM (Invitrogen™) and 4 μM Ethidum homodimer-1 (Invitrogen™) in PBS for 30 min at 37°C. Following staining, hydrogels were rinsed with PBS, and imaged using a Zeiss LSM confocal microscope (Zeiss Canada) with a 10X objective. A minimum of 5 hydrogels were imaged at each timepoint (n=5).

### 5.8 In vitro bioluminescence imaging

The luminescence signals of transduced hASCs in the MGC-RGD+PEG(PTMC-A)_2_ hydrogels were assessed following encapsulation, as well as following 24 h, 3 d, 7 d, and 14 d of culture *in vitro*. At each timepoint, hydrogels were transferred into black bottom 96-well plates for the luminescence readings, and incubated in 100 μL D-luciferin (Syd Labs) in PBS at a concentration of 1.25 mg/mL. Luminescence imaging was performed using an IVIS Lumina XRMS Imaging System (PerkinElmer), with images captured using auto-exposure time until a peak average radiance had been reached. Photographic images were used to draw an ROI surrounding the hydrogels, to quantify average radiance (p/s/cm^2^/sr). Four hydrogels were assessed at each time point (n=4).

### 5.9 Femoral artery ligation-induced critical limb ischemia (FAL-CLI) model

All animal procedures were performed in accordance with the rules and regulations set by the Canadian Council on Animal Care and were approved by the Animal Care Committee at Western University (Animal Use Protocol #2019-024). *In vivo* studies were performed using 8-week old athymic *nu/nu* mice (Crl:NU-Foxn1nu, Charles River). Unilateral hindlimb ischemia was induced in the mice through ligation and excision of a 2-5 mm portion of the femoral artery and vein in the right hindlimb, following established protocols [36], [37]. Anesthesia was induced and maintained using inhaled isoflurane.

Mice were allowed to recover from the FAL surgery for 24 h, and then randomly assigned to the various treatment groups. Four treatment groups were assessed: (i) hASCs in MGC-RGD+PEG(PTMC-A)_2_ hydrogels, (ii) hASCs in saline, (iii) MGC-RGD+PEG(PTMC-A)_2_ hydrogel alone controls, and (iv) saline alone controls. For all treatment groups, a total volume of 20 μL was injected intramuscularly into the surgical limb, just distal to the site of ligation, while the mice were under anesthesia. Polymer solutions were prepared as described above for *in vitro* encapsulation. hASCs were delivered at a concentration of 2x10^7^ cells/mL (4x10^5^ cells/injection). The study was performed using the transduced hASCs with n=6 mice/hASC treatment group, and n=3 mice/control treatment group.

### 5.10 Laser Doppler perfusion imaging assessment of limb perfusion

LDPI was used to assess limb perfusion at days 1 (prior to treatment), 4, 7, 14, 21, 28, and 35. Mice were anesthetized with isoflurane and warmed at 37°C on a heating pad for 5 min. The mice, maintained under isoflurane anesthesia, were imaged in a supine position using a Moor LDI-2 laser Doppler imaging system (Moor Instruments). The average flux in the ischemic (surgical) and control limbs were quantified within a ROI encompassing the foot and ankle joint. The perfusion ratio (PR) was calculated as the ratio between the average flux in the surgical and control limbs and normalized relative to the perfusion ratio at day 1. All mice included in the study had a day 1 PR <0.15.

### 5.11 In vivo bioluminescence imaging assessment of cell retention

*In vivo* BLI was performed to assess the retention of transduced hASCs within the ischemic limbs in the FAL-CLI model at days 1 (following injection), 4, 7, 14, 21, 28, and 35. Mice were anaesthetized using isoflurane and received an intraperitoneal injection of 100 μL of 30 mg/mL D-luciferin (Syd Labs), and subsequently imaged using an IVIS Lumina XRMS *In Vivo* Imaging System (PerkinElmer). An ROI of consistent size was drawn around the site of injection and used to quantify the average radiance (p/s/cm^2^/sr) of the luminescence signal. Images were taken using auto-exposure until the average radiance reached a peak value. Average radiance values were normalized to the day 1 values for each mouse.

### 5.12 Immunofluorescence analysis of immune cell recruitment

The athymic *nu/nu* mice from the first two cohorts were euthanized 35 days following treatment. The mice from the third cohort were euthanized 12 days following treatment for the hASCs in MGC-RGD+PEG(PTMC-A)_2_ group or 14 days following treatment for all other groups due to facility access restrictions resulting from the COVID pandemic. The skin covering the hindlimbs was removed, and the entire thigh muscle was harvested and fixed for 24 h in 10% neutral buffered formalin solution (Fisher Scientific). Samples were then subjected to a sucrose gradient (10%, 20%, 30% sucrose in PBS; incubated for 24 h in each solution), and embedded in frozen section compound (VWR), frozen on dry ice, and stored at -80°C until further processing.

Samples were cryo-sectioned at a thickness of 10 μm and stained to assess host immune cell recruitment with antibodies for murine CD45. Sections were fixed in acetone (Fisher Scientific) for 10 min at -20°C, followed by three 2 min washes in PBS. Samples were then blocked in a solution of PBS with 0.05% tween (Bio-shop) and 10% donkey serum for 1 h at room temperature. Staining for CD45 (1:100 in blocking solution, goat polyclonal anti-CD45, Bio-Techne®, AF114) was performed overnight at 4°C. Sections were washed 3 X 2 min in PBS, and then incubated with the secondary antibody donkey anti-goat IgG Alexa Fluor™ 546 (1:400 in PBS with 0.05% tween, A11056, Invitrogen) for 1 h at room temperature. Sections were washed 3 X 2 min in PBS, then mounted using mounting medium with DAPI (Abcam) to stain cell nuclei. Stained cross sections at 3 depths at least 200 μm apart were imaged per mouse using a Zeiss Imager M2 Microscope (Zeiss Canada) with a 10X objective. Controls without primary antibody were included to confirm staining specificity. Semi-quantitative scoring of areas of dense CD45^+^ staining was performed. The tissue sections were given scores of 0 (no dense CD45^+^ staining outside of the ligated vessel region), 1 (only one small region of dense CD45^+^ staining), 2 (one medium region of dense CD45^+^ staining or multiple small regions) or 3 (at least one large region of dense CD45^+^ staining). Scoring was performed in a blinded manner.

### 5.13 Statistical Methods

All data was presented as the group mean ± standard deviation, unless otherwise noted. All statistical analyses were performed using GraphPad Prism 9 software. Area under the curve (AUC) analyses were performed on the normalized *in vivo* LDPI and BLI data [38]–[41]. Differences in the immunophenotype and viability of transduced and non-transduced hASCs were assessed using multiple paired t-tests (paired across hASC donors), with Holm-Šídák post-hoc correction. Differences in *in vitro* luminescence values across timepoints were assessed using one-way ANOVA with Tukey’s post-hoc test for multiple comparisons. Differences in perfusion ratio data between treatment groups at each time point were assessed using two-way repeated measures (RM) ANOVA, with Tukey’s post-hoc test for multiple comparisons. Area under the curve data for perfusion ratio was assessed using one-way ANOVA with Tukey’s post-hoc test for multiple comparisons. Differences between treatment groups at each timepoint for the *in vivo* BLI data were assessed using multiple unpaired t-tests with a false discovery rate set to 5%, while differences in the AUC for the same data were assessed using an unpaired t-test. Data was considered statistically significant when p < 0.05.

## 6 Acknowledgements

Operational funding for this study was provided by the Heart and Stroke Foundation of Canada (Grant-in-Aid, G-19-0026269) and the Canadian Institutes of Health Research (CIHR MOP #378189). Fiona Serack was supported by an Ontario Graduate Scholarship and a Transdisciplinary Bone & Joint Training Award from the Bone and Joint Institute at Western University. Drs. A. Grant, R. Richards, and D. Matic are acknowledged for their clinical collaborations in providing the adipose tissue samples, as well as Drs. Pascal Morissette Martin, Amanda Hamilton, and Ying Xia for their technical support. Additionally, the authors would like to thank the London Regional Flow Cytometry Facility for access to flow cytometry equipment.

## 7 Authorship Contribution Statement

Dr. Fiona E. Serack performed the experiments and data analysis, prepared the figures, and drafted the manuscript. Dr. Lauren E. Flynn contributed to the design and direction of the study, and provided critical feedback on data analysis and interpretation, as well as contributed to manuscript writing. In addition, Dr. John Ronald provided scientific guidance with the lentiviral transduction and access to the *in vivo* imaging system platform. Finally, Dr. Brian G. Amsden and Dr. David A. Hess provided guidance on polymer synthesis and the FAL-CLI model, respectively, and assisted with manuscript editing.

## 8 Author Disclosure Statement

The authors declare that there are no potential conflicts of interest associated with this research.

## 9 Funding Statement

Funding for this study was provided by the Heart and Stroke Foundation of Canada (Grant-in-Aid, G-19-0026269) and the Canadian Institutes of Health Research (CIHR MOP #378189).

## Notes

### Competing Interest Statement

The authors have declared no competing interest.

### Summary of Updates

Pilot NOD/SCID in vivo data removed (original Figure 2). Format updated for resubmission.

