## Supplementary Figure S2 for "Delivery of human adipose-derived stromal cells within *in situ* forming chitosan/PEG-PTMC hydrogels induces adverse outcomes in a femoral artery ligation model in athymic *nu/nu* mice"

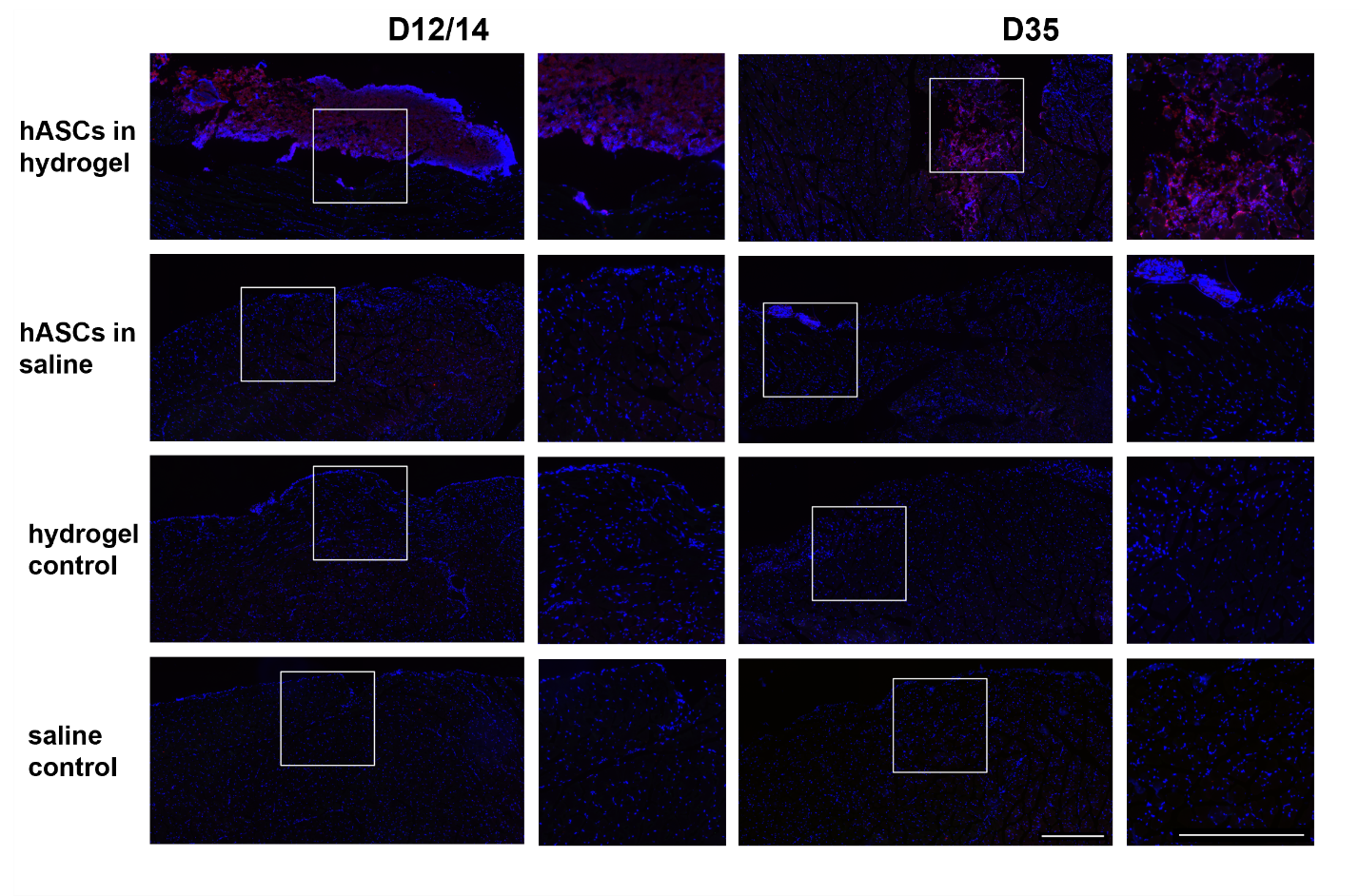

Supplementary Figure S2. CD45^+^ staining results showing a more potent inflammatory response within the ischemic muscle in a higher fraction of the mice that received hASCs in the hydrogel. Representative IHC images of mouse thighs harvested post-euthanasia at days 12/14 or 35 following FAL (DAPI^+^ cells in blue and CD45^+^ cells in red). Images (10X) were taken across the entire cross-section of the muscles and tiled. Scale bars represent 500 μm.
