## Supplementary Figure S1 for "Delivery of human adipose-derived stromal cells within *in situ* forming chitosan/PEG-PTMC hydrogels induces adverse outcomes in a femoral artery ligation model in athymic *nu/nu* mice"

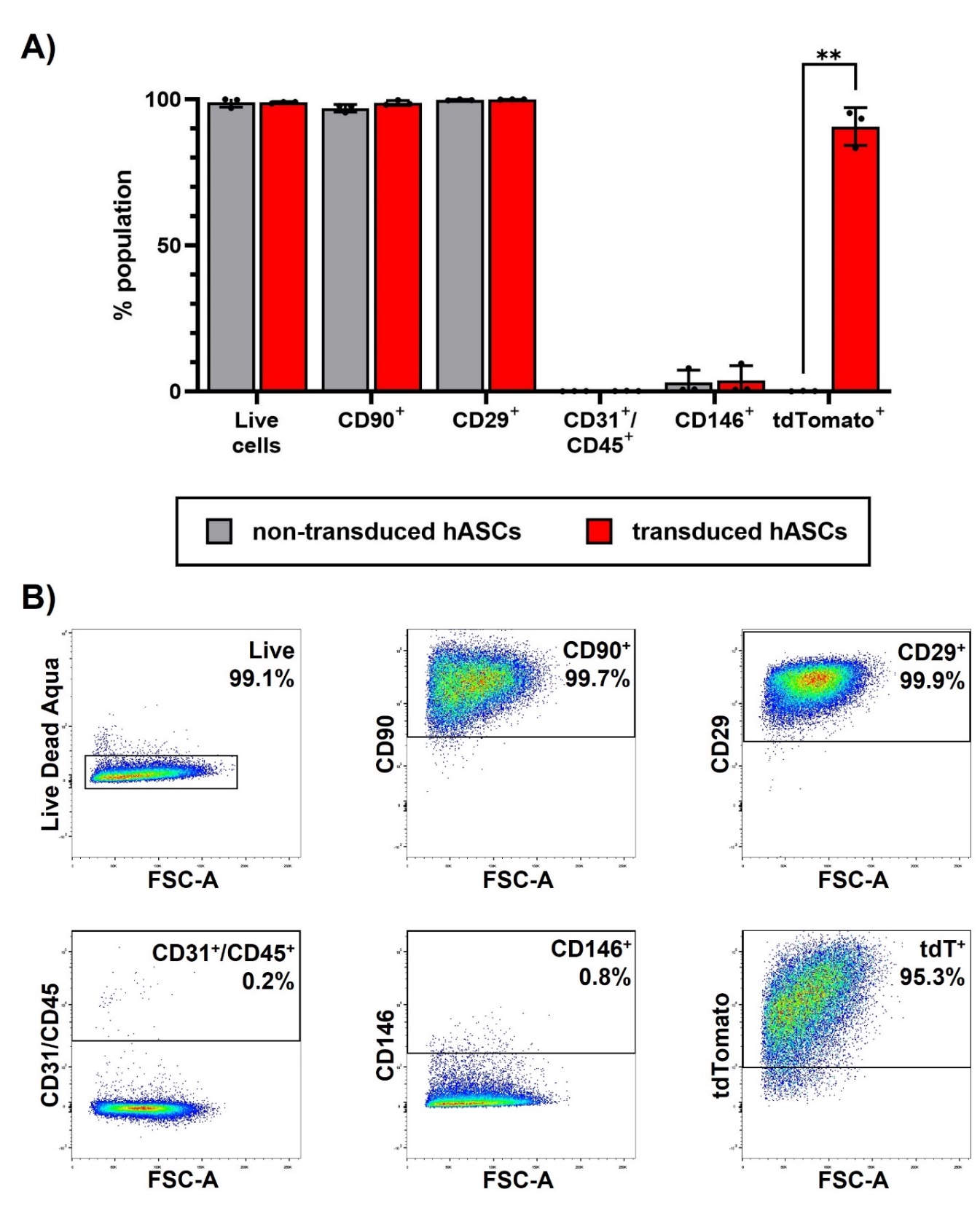
**Supplementary Figure S1. Transduction of hASCs did not alter cell viability or surface marker expression patterns.** (A) Flow cytometry was used to characterize the hASC viability and immunophenotype. Live cells were analyzed by the expression of the mesenchymal markers CD90 and CD29, the endothelial and leukocyte markers CD31 and CD45 respectively (placed on the same channel), the pericyte marker (CD146), and tdTomato. The immunophenotype of the hASCs was as expected, and no differences were observed between the transduced and non-transduced cells, except for in the expression of tdTomato (** p<0.01). Data presented as mean ± SD. Differences between transduced and non-transduced cells were detected using multiple paired t-tests, with Holm-Šídák post-hoc correction (N = 3 different hASC donors). (B) Representative flow cytometry plots showing hASC immunophenotype on transduced hASCs. Cells were first gated based on size and singlets (not shown), then gated based on viability. Live cells were then gated for the cell surface markers, or tdTomato based on FMO controls.
