## Supplementary Figure S3 for "Delivery of human adipose-derived stromal cells within *in situ* forming chitosan/PEG-PTMC hydrogels induces adverse outcomes in a femoral artery ligation model in athymic *nu/nu* mice"

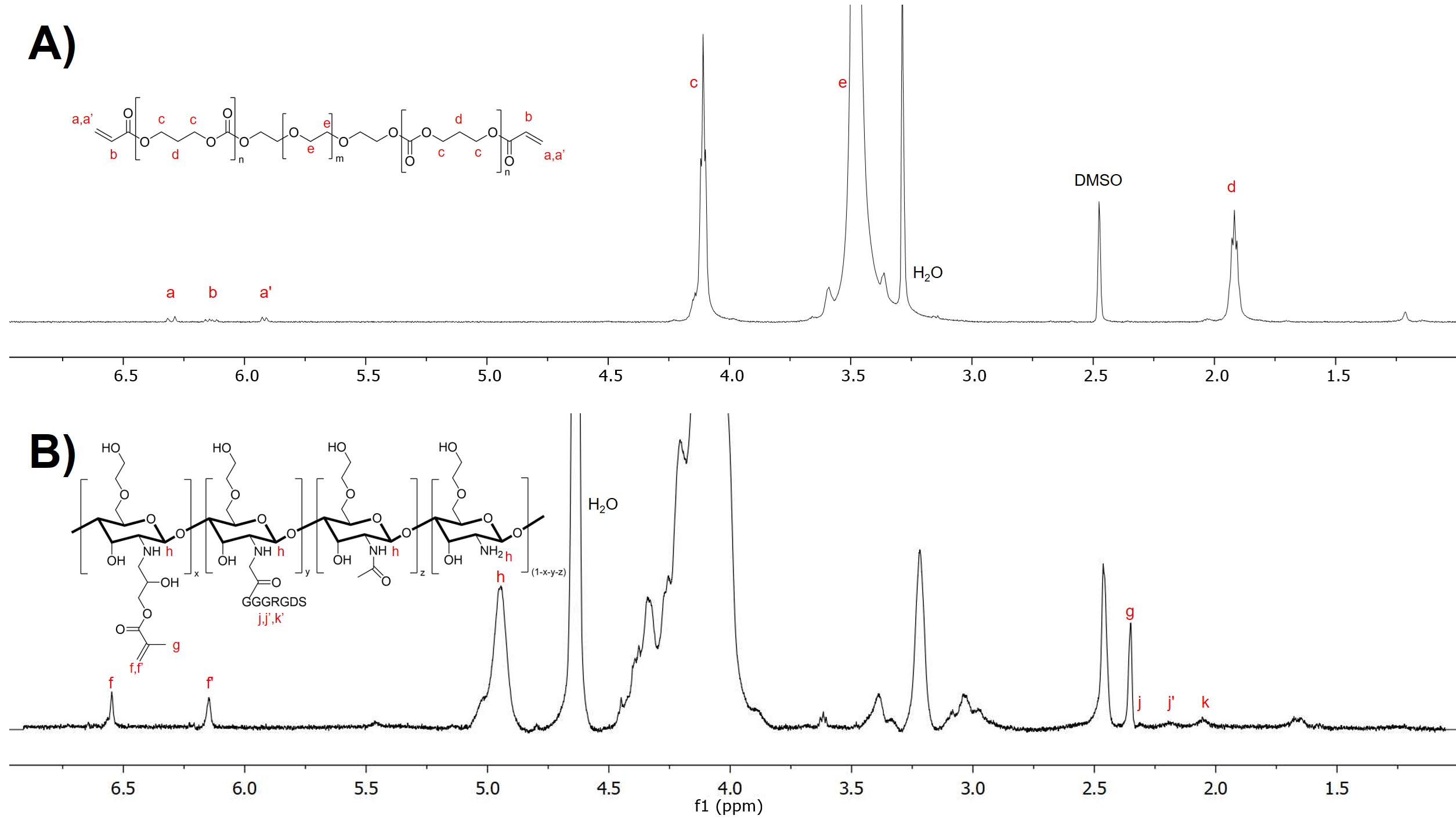

Supplementary Figure S3. ^1^H NMR was used to confirm the structure and degree of functionalization of PEG(PTMC-A)_2_ (A) and MGC-RGD (B). (A) Molecular ratios of PEG_20_(PTMC_2_-A)_2_ were determined, and the degree of acrylation was shown to be 85%, based on the ^1^H NMR spectrum. (B) For MGC-RGD, the degree of methacrylation was 5%, and the degree of RGD functionalization was 5%, as calculated from the ^1^H NMR spectrum. The fraction of residual acetyl groups was reported by the supplier as 15%.
